# Common viral infections inhibit egg laying in honey bee queens and are linked to premature supersedure

**DOI:** 10.1101/2024.04.16.589807

**Authors:** Abigail Chapman, Alison McAfee, David R. Tarpy, Julia Fine, Zoe Rempel, Kira Peters, Rob Currie, Leonard J. Foster

## Abstract

With their long lives and extreme reproductive output, social insect queens have escaped the classic trade-off between fecundity and lifespan but evidence for a trade-off between fecundity and immunity has been inconclusive. This is in part because pathogenic effects are seldom decoupled from effects of immune induction. We conducted parallel, blind virus infection experiments in the laboratory and in the field to interrogate the idea of a reproductive immunity trade-off and better understand how these ubiquitous honey bee stressors affect queen health. We found that queens injected with infectious virus had smaller ovaries and were less likely to recommence egg-laying than controls, while queens injected with UV-inactivated virus displayed an intermediate phenotype. In the field, heavily infected queens had smaller ovaries and infection was a meaningful predictor of whether supersedure cells were observed in the colony. Immune responses in queens receiving live virus were similar to queens receiving inactivated virus, and several of the same immune proteins were negatively associated with ovary mass in the field. This work solidifies the relationship between virus infection and symptoms associated with queen failure and suggests that a reproductive-immunity trade-off is partially, but not wholly responsible for these effects.

## Introduction

As the sole reproductive female in a colony of thousands, honey bee (*Apis mellifera*) queens are the cornerstones of the colonies that they head. Understanding what factors impact their health and reproductive potential is therefore of great interest to the beekeeping industry, which provides pollination services valued annually in the billions globally (Gallai et al., 2009; Khalifa et al., 2021). While honey bee queens can live for 5 years or more (Winston, 1987), beekeepers regularly replace queens at much younger ages, often after less than 1 year (Amiri et al., 2017). It is still unclear, though, exactly what factors and in what combinations contribute to the symptoms of “queen problems” or poor reproductive output, which are regularly reported as one of the main perceived causes of colony loss (Canadian Association of Professional Apiculturists (CAPA), 2024; Seitz et al., 2022; vanEngelsdorp et al., 2008).

Honey bee queens mate over a short period of time with multiple drones at a few weeks of age and store the sperm they acquire for the rest of their lives. The remaining quality and quantity of this stored sperm is one determining factor of queen quality – if a queen lacks viable sperm with which to fertilize eggs, either because she did not obtain enough through mating or because the remaining sperm is no longer viable, her productivity will decline (Pettis et al., 2016; Tarpy & Olivarez Jr., 2014). Sperm quality can be affected by abiotic factors, such as temperature stress (McAfee et al., 2020; Pettis et al., 2016) and pesticide exposure (Chaimanee et al., 2016). Additionally, the age of larvae that beekeepers graft when rearing new queens can impact the physiology and size of queens (Woyke, 1971), with older larvae developing into worker-like queens, which impacts their future reproductive potential (De Souza et al., 2019; Dedej et al., 1998; Rangel et al., 2013; Tarpy & Mayer, 2009). Queens perceived as failing (exhibiting drone laying in worker cells, or producing poor brood patterns) by beekeepers have been shown to have reduced sperm viability (Pettis et al., 2016) and we previously reported that they also have smaller ovaries compared to their healthy counterparts of similar age and in similar environments at the same time of year (McAfee et al., 2021). In the same study, we also linked natural viral infection to queen problems in field operations, finding that beekeeper-identified failing queens had higher viral titres and that queens with higher viral titres, regardless of their performance status (healthy or failing), also had smaller ovaries (McAfee et al., 2021). We later established the causal rather than correlative nature of this relationship wherein injection with Israeli acute paralysis virus (IAPV) caused a reduction in ovary mass—the same phenotype observed in queens identified as “failing” from the field (Chapman et al., 2022). However, experiments investigating changes in actual reproductive output in response to viral infection, particularly with the largely asymptomatic infections commonly encountered in the field, have not been conducted.

Many of the honey bee viruses commonly found in colonies can infect queens, but their pathological effects are not yet fully defined. Deformed wing virus (DWV), has two common genotypes: A and B, the latter of which was detected more recently, is more virulent and is potentially replacing genotype A (Paxton et al., 2022; Tehel et al., 2019). This virus is commonly vectored by the parasitic mite, *Varroa destructor*, but is also transmitted horizontally, vertically, and sexually (de Miranda & Fries, 2008; de Miranda & Genersch, 2010; Ravoet et al., 2015). Queens rarely show any overt symptoms of infection, and the deformed wing phenotype seen in workers has only been reported a single time in a queen (Williams et al., 2009). However, the virus has been detected in all queen tissues, including the head, thorax, gut, fat body, ovaries, and spermatheca (Amiri et al., 2016; Francis et al., 2013). High levels of DWV infection have been associated with ovarian degeneration and egg-laying deficiencies in queens (Gauthier et al., 2011), but few other studies have examined the pathological effects of DWV infection on queens, with most focusing solely on detection of the virus in various organs. Other viruses, including sacbrood virus (SBV), black queen cell virus (BQCV), acute bee paralysis virus (ABPV), and IAPV have also been detected in various queen tissues including the ovaries, but generally only in low titers, and the effects on queen health have been even less well-studied (Amiri et al., 2017, 2020; Chen et al., 2005; Francis et al., 2013). A recent study on the prevalence of a newly identified honey bee virus, Solinvivirus-1, found that apiaries testing positive for the virus were nearly twice as likely to contain queenless colonies (Ryabov et al., 2023), and our previous findings linking viral infection to queen failure suggests that, although infections are often hidden in queens, they may still be compromising queen health and reproductive output.

Determining the mechanism of this effect is also biologically intriguing because, while the pathogenic effects of a viral infection could be directly compromising a queen’s health and reproductive output, a reproductive-immunity trade-off may also be at play. A compromise between reproduction and immunity has been well studied in many insects (Schwenke et al., 2016), but not previously in honey bees. This compromise, which is generally considered to be driven by allocation of limited resources (since both reproduction and immune activation are energetically costly) could be particularly relevant for honey bee queens, who can sustain an enormous reproductive output and can lay up to their own bodyweight in eggs every day (Rueppell et al., 2016). It is reasonable to assume that sustaining such an energetically intensive process would come with costs, not only to immune function, but to longevity and somatic maintenance in general, as is the case in many insects—many of which are not as specialized for reproduction as the honey bee queen.

However, honey bees and other highly social insects (ants and termites) have escaped the trade-off with lifespan, and exhibit both enormous reproductive output and increased longevity compared to the non-reproductive castes (Blacher et al., 2017; Remolina & Hughes, 2008; Rueppell et al., 2016; von Wyschetzki et al., 2015). This uncoupling of fecundity and longevity appears to be due in part to a rewiring of the endocrine network which signals resources to be directed towards one process or the other in solitary insects (Corona et al., 2007; Remolina & Hughes, 2008; Rodrigues & Flatt, 2016) and involves insulin/insulin-like growth factor (IIS), juvenile hormone (JH), and vitellogenin (Vg). Given that social insects have escaped at least one compromise, it therefore calls into question whether they may have also evaded the trade-off with immunocompetence. However, there is conflicting evidence for other social insects. Black garden ant (*Lasius niger*) queens appear to have removed the main mediator of a trade-off by uncoupling JH from its role as stimulator of reproduction (as is the case in solitary insects) while maintaining its role as a suppressor of immune activation (Pamminger et al., 2016). Conversely, a different study in ant queens found evidence for a trade-off after wounding, in which the queens exhibited reduced egg laying with an increase in immune expression and a decrease in germ cell development genes (von Wyschetzki et al., 2016). In social bee species, the limited evidence so far is similarly conflicting. We have recently suggested that in bumble bees, what appears as evidence for a trade-off between immunity and reproduction in the protein expression patterns of queens may actually be due instead to differing selection pressures at different stages of a queen’s life cycle, which cannot be de-coupled from her reproductive activation (McAfee et al., 2024). In honey bees, it has been shown that reproductive potential does not reduce the heat-shock response (a response which includes antiviral immune effectors, McMenamin et al., 2020), which is unique among the invertebrates that have been studied to date (Shih et al., 2020). In contrast, we have previously found evidence for a trade-off in honey bee queens: sperm viability, another important reproductive metric, is negatively correlated with lysozyme, a conserved immune effector (McAfee et al., 2021), and injection with IAPV caused a decrease in the transcriptional expression of vitellogenin and an increase of heat shock protein expression (Chapman et al., 2022). We are interested in determining if indeed a trade-off is driving the effects of viral infection we observe because, if so, this means that even the ubiquitous levels of virus infection present in colonies—which may not be “making queens sick” or causing overt disease —could still be reducing the reproductive success of a queen.

Our previous experimental infections with IAPV were conducted over a short time frame (3 days) and measured only ovary mass to quantify the impact to reproduction. However, poor queen performance is a chronic symptom displayed over time, not an acute effect. In this study, we therefore aimed to determine if the effects of experimental virus infection extend to an observable effect on actual reproductive output, beyond just smaller ovaries, by employing more realistic experimental settings and a longer timeframe. We conducted two separate experiments in parallel in which we observed queens experimentally infected with a combination of BQCV and DWV-B either in laboratory cages (queen monitoring cages, or QMCs) (Fine et al., 2018) which allow queens to continue laying eggs for one week following infection, or in colonies for 6 weeks following infection. To address the potential presence of a reproductive-immunity trade-off, we also injected a group of queens in the cage experiment with a UV-inactivated version of the same viral inoculum to decouple the role of immune activation from viral pathogenesis.

## Results

### Cage experiment - virus titres

Young, mated honey bee queens were injected with a) a combination of infectious BQCV and DWV-B, b) the same mixture of virus which had been inactivated with ultraviolet light, or c) saline (injection control) and subsequently kept in incubated cages which enabled them to lay eggs for 7 days following injection. Because queens are likely to have background infections, we assessed the titres of common viruses (BQCV, DWV-A, DWV-B, and SBV) in all groups using RT-qPCR. As expected, there were no differences in viral titres in the inactivated group compared to the saline-injected control, and in the infected group DWV-B was increased and BQCV was marginally increased (Figure 1a). Correspondingly, the infected group had a significantly higher level of “total virus,” a metric representing the sum of the titres of the individual viruses (Figure 1b), which we have previously determined to be an important predictor of ovary mass in queens collected from the field than the levels of any singular virus (Chapman et al., 2022). See Table 1 for complete statistical details.

**Figure 1.**
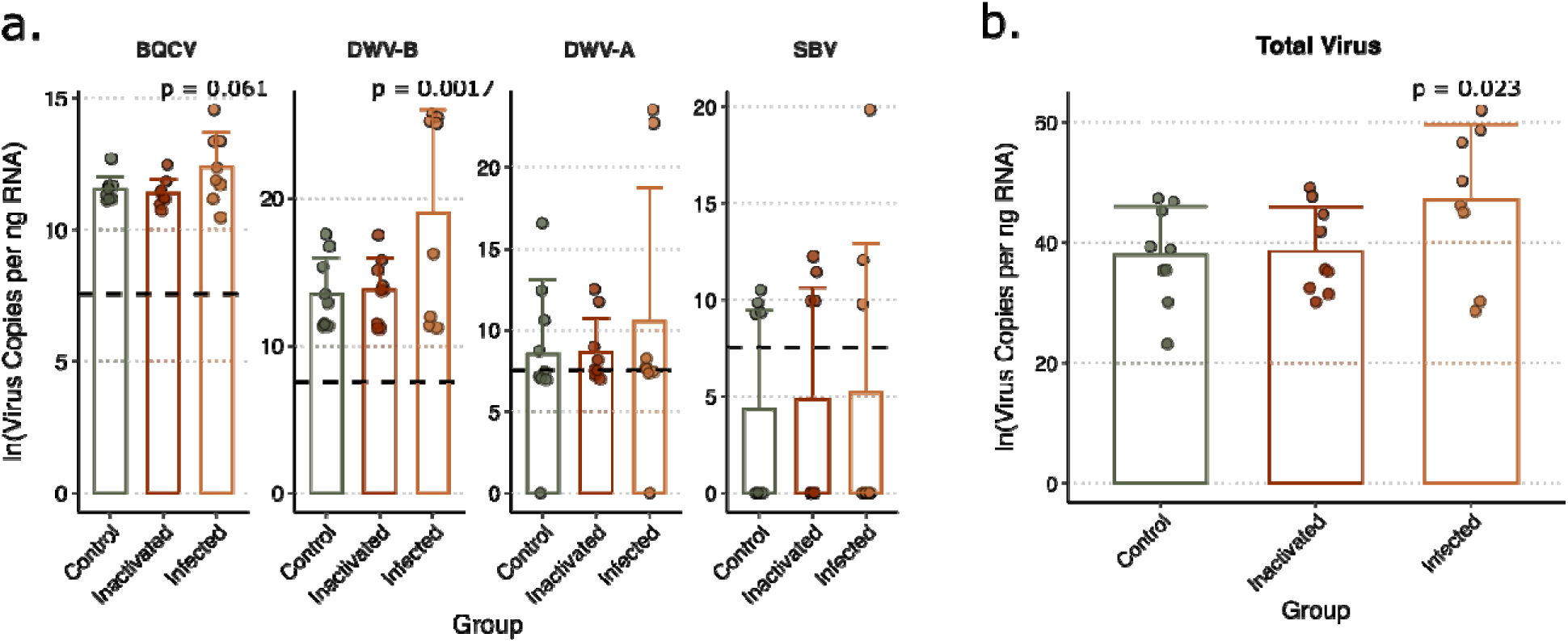
Virus titres of cage experiment queens. Bars show mean values and error bars represent standard deviation; p-values are shown relative to the control group. Control queens were injected with saline, infected queens were injected with a combination of BQCV and DWV-B, and queens in the inactivated group were injected with the same combination of virus that was UV-inactivated (*n* = 9 for all groups). Queens were kept in quee monitoring cages in an incubator with attendants and sacrificed 7 days after injection. **a.** Titres of BQCV and DWV-B were higher in the infected group, while there were no differences in the titres of DWV-A and SBV between groups. The dashed horizontal lines represent the RT-qPCR quantification threshold. **b.** The summed total virus titre was higher in the infected group relative to the control group.

**Table 1.** Statistical parameters for cage experiment*.

| Response | Fixed effects | Model | Estimate | df<sup>b</sup> | \|t/z\| | p |
| --- | --- | --- | --- | --- | --- | --- |
| BQCV | Group (control group as reference) | Linear mixed model (lmer)<sup>a</sup> | Live: 0.81 | 23 | 1.97 | 0.061 |
|  |  |  | Inactive: -0.16 |  | -0.40 | 0.69 |
| SBV |  |  | Live: 0.88 | 23 | 0.29 | 0.77 |
|  |  |  | Inactive: 0.51 |  | 0.17 | 0.86 |
| DWV-B |  |  | Live: 5.12 | 21 | 3.59 | 0.0017 |
|  |  |  | Inactive: -0.37 |  | -0.27 | 0.79 |
| DWV-A |  |  | Live: 1.75 | 21 | 1.045 | 0.31 |
|  |  |  | Inactive: 0.071 |  | 0.044 | 0.97 |
| Total virus |  |  | Live: 8.75 | 21 | 2.45 | 0.023 |
|  |  |  | Inactive: 0.37 |  | 0.11 | 0.92 |
| Ovary mass | Group | Linear mixed model (lmer)<sup>a</sup> | Live: -6.74 | 22 | -3.31 | 0.0032 |
|  |  |  | Inactive: -3.61 |  | -1.77 | 0.090 |
| Ovary mass | BQCV | Linear mixed model (lmer)<sup>a</sup> | -2.32 | 22 | -2.28 | 0.033 |
|  | DWV-B |  | -0.56 | 24 | -2.72 | 0.012 |
|  | DWV-A |  | -0.062 | 10 | -0.24 | 0.82 |
|  | SBV |  | -0.21 | 22 | -1.22 | 0.23 |
|  | Total virus |  | -0.26 | 15 | -2.62 | 0.020 |
| Recommence laying eggs after injection? (Y/N) | Group (control group as reference) | Generalized linear model, binomial (GLM) | Live: -3.33 | n/a<sup>c</sup> | -2.51 | 0.012 |
|  |  |  | Inactive: -0.83 |  | -0.62 | 0.53 |
| # eggs laid | Total virus | Linear mixed model (lmer)<sup>a</sup> | -2.51 | 24 | -2.64 | 0.014 |

\*N = 27; n = 9 per group. All virus titres (virus copies per ng RNA) were transformed by taking the natural logarithm
<sup>a</sup>Experimental batch was included as a random effect
<sup>b</sup>For linear mixed models, degrees of freedom were estimated using Satterthwaite's method.
<sup>c</sup>Generalized linear models do not report degrees of freedom; for this model there were 27 observations and 3 groups.

### Cage experiment – fertility analysis

The ovary masses of queens injected with live virus significantly decreased after 7 days relative to the saline-injected queens (linear mixed model with batch as a random effect, t = −3.31, p = 0.0032) (Figure 2a), which is consistent with what we previously observed after injection with IAPV (Chapman et al., 2022). The ovary masses of queens injected with the inactivated virus also tend to be non-significantly reduced relative to the control, though the effect size is about half that of the live virus group (linear mixed model with batch as random effect, t = −1.77, p = 0.090). Regardless of group, ovary mass was negatively associated with levels of BQCV, DWV-B, and total virus load (Figure 2b, see Table 1 for statistical reporting).

**Figure 2.**
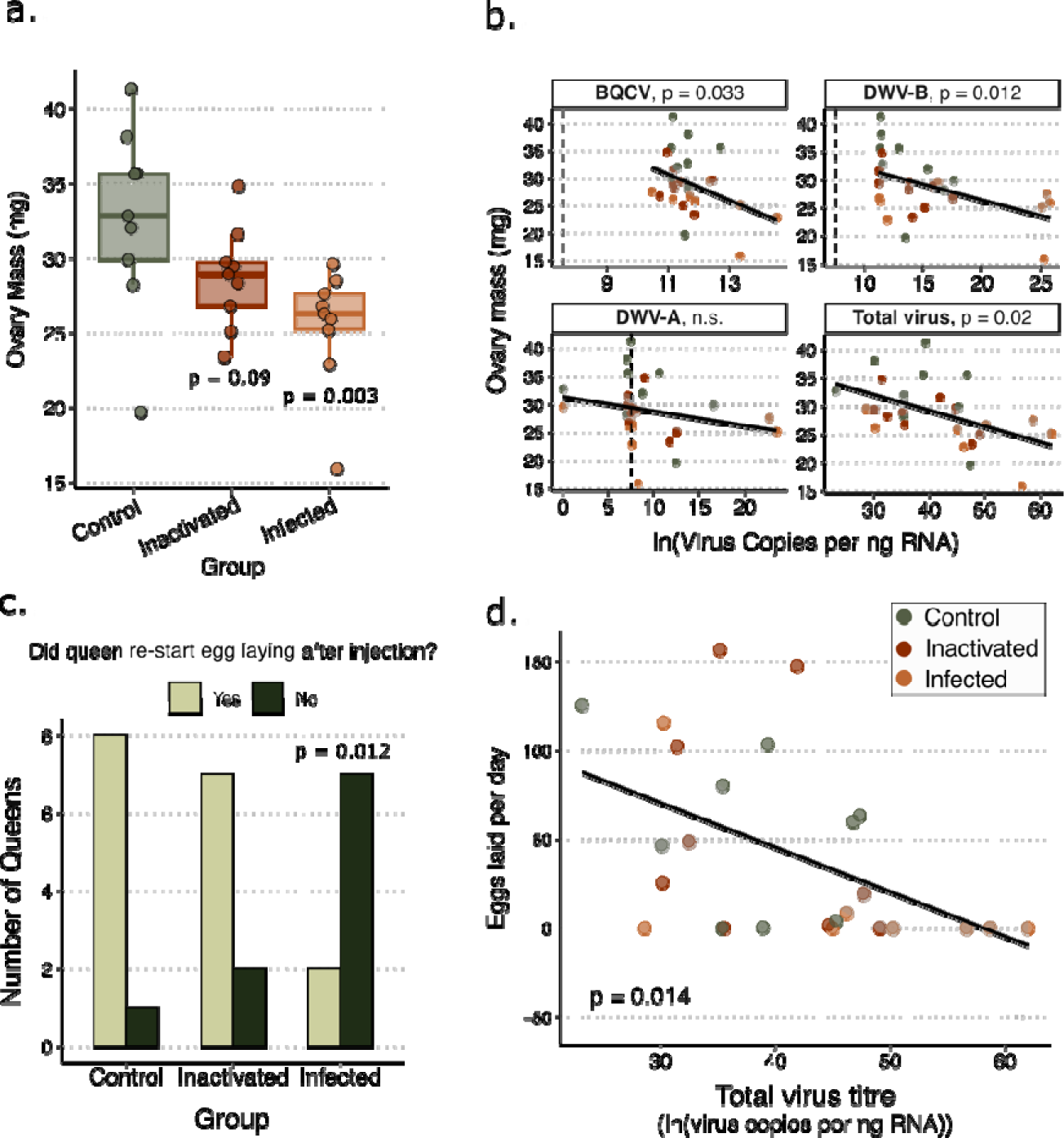
Viral infections reduced ovary mass and egg laying in cage experiment queens. **a.** Ovary mass decrease significantly in queens injected with live virus relative to the control group, with a non-significant effect observed in queens injected with inactivated virus. **b.** Ovary mass is negatively associated with levels of BQCV, DWV-B, and total virus load. The dashed vertical lines represent the RT-qPCR quantification threshold. **c.** Queens injected wit live virus were significantly less likely to recommence egg laying 7 days after injection compared to control queens. **d.** The number of eggs laid per day, averaged over days 5-7 after injection, was negatively associated with the total virus titre across all queens. See Table 1 for detailed statistical reporting.

Queens were kept in specialized queen monitoring cages (QMCs) which enabled us to record their egg laying. All queens were confirmed to be laying consistently before being injected, but queens injected with live virus were 85% less likely to recommence laying in the 7 days following injection compared to the control queens (binomial generalized linear model, estimate = −3.33, z = −2.51, p = 0.012), as shown in Figure 2c. Mirroring the results of ovary mass, regardless of group, for every additional unit of total virus infection queens laid approximately 2.5 fewer eggs per day over the final 3 days before being sacrificed (Figure 2d, linear model, t = −2.64, p = 0.014).

### Cage experiment – hemolymph proteome analysis

To investigate how the immune response of queens injected with UV-inactivated virus differed from those infected with live virus, we performed quantitative proteomics on the hemolymph of queens. Of the 1,071 identified protein groups (1% FDR) and 933 protein groups quantified in at least 75% of samples, 23 proteins were differentially expressed in the live infected group, and 15 were differentially expressed in the UV-inactivated group relative to the control (Figure 3, 5% FDR, Benjamini-Hochberg correction). Between these two groups, 12 proteins are shared, including many which are known to be associated with immunity. IRP30, for example, is a secreted immune protein found in a wide variety of hymenopteran species and is co-expressed with carboxylesterase (Albert et al., 2011; Dong et al., 2020), which is also upregulated in both groups. Transferrin mediates iron-sequestration as an innate immune strategy to limit the availability of the element for acquisition by pathogens (Iatsenko et al., 2020). Beta-1,3-glucan binding protein is a pattern-recognition receptor in invertebrates for the glucans found on fungal cell walls, which triggers the phenoloxidase cascade, possibly by interacting with a serine protease (Ma & Kanost, 2000; Vargas-Albores & Yepiz-Plascencia, 2000). Additionally, phenoloxidase activating factor, a modular serine protease, and several of the antimicrobial peptides including defensin, hymenoptaecin, and apidaecin are all upregulated. The only immune effector not shared between groups, lysozyme, was significant in only the group receiving inactivated innoculum. A channel protein from *Vairimorpha apis* (formerly *Nosema apis* (Tokarev et al., 2020)) was also upregulated in both the inactivated and infected groups.

**Figure 3.**
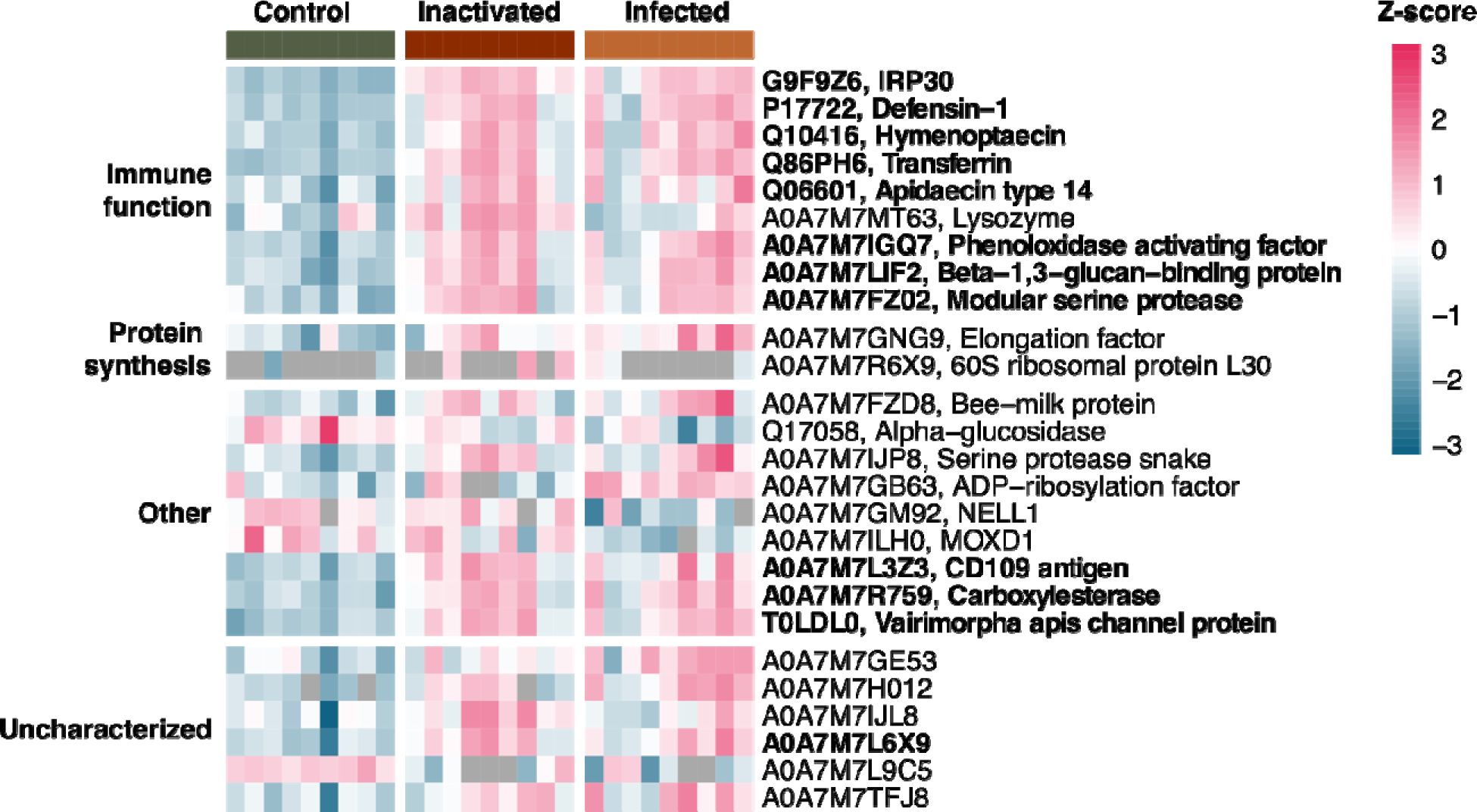
Hemolymph immune proteins were differentially expressed between groups in cage experiment queens. Each row represents a protein, and each column represents a sample, showing the Uniprot accessio number. Proteins that were not quantified in a sample (“NA”) are shown as grey tiles. Proteins in bold were significant in both control-inactivated and control-infected contrasts (5% FDR, Benjamini-Hochberg correction).

### Cage experiment – perturbation caused by the cage environment

For experiments in which bees are kept in the laboratory, it is often unknown exactly how disruptive the laboratory system is to their natural functioning. To address this, we sacrificed three “baseline” queens from each of the three batches we received for the cage experiment immediately upon arrival for comparison to the control queens which were subsequently kept in the QMCs. The ovaries of the baseline queens were significantly larger than the ovaries of control queens that were sacrificed 2 weeks later (Supplementary Figure 1a). This is unsurprising, given that the baseline queens had been actively laying eggs in colonies only ∼24 hours before being sacrificed, while the control queens had been kept in cages which, while allowing them to lay, did restrict their laying compared to a colony environment. To assess the impact of the QMCs more thoroughly, we also performed proteomics on the hemolymph of the baseline queens and found that even though 39 proteins in the hemolymph were differentially expressed in the baseline queens compared to the controls (5% FDR, Benjamini-Hochberg correction), only 3 proteins were overlapping with the proteins differentially expressed either of the experimental inactivated or infected groups (Supplementary Figure 1b). This indicates that even though the cage environment clearly has an impact on the physiology of queens over this two-week period, those effects do not appear to overlap substantially with the effects we observed as a result of our experimental injections.

### Field experiment – virus titres and mite levels

For the field experiment, queens were injected with either saline or a combination of BQCV and DWV-B in the same manner as the cage experiment, placed in colonies, and assessed weekly for 7 weeks following injection. We again assessed the virus loads of the queens at the end of the experiment using RT-qPCR. As shown in Figure 4a, BQCV was higher in the infected group while DWV-B was actually lower in the infected group (see Table 2 for statistical details). DWV-A was marginally higher in the infected group and SBV was only detected in two queens. Correspondingly, there was no significant difference in the total virus level between the two groups. We also randomly sampled workers from brood frames at both the beginning and the end of the experiment to assess the levels of viruses present in the colony. The viral load in workers at the beginning of the experiment was a significant predictor of the total weight gained by the colony over the course of the experiment (Figure 4b) while the queen viral load was not. Interestingly, we found that the level of DWV-B in queens was significantly linked to the titres of DWV-B present in the workers at the end of the experiment, though there was no correlation between queen titres and worker titres for the other viral metrics after controlling for the effect of site (Figure 4c). This trend with DWV-B appears to be mainly driven by colonies in the control group. We also examined the correlations of worker viral titres, queen viral titres, and the levels of mites in the colonies at the end of the experiment to gain a better understanding of how these metrics covaried. While mite load is positively correlated with levels of both variants of DWV in both queens and workers, the correlation is not significant after adjusting for multiple hypothesis testing (Bonferroni) (Figure 4d). There were no differences in mite loads between the experimental groups and colonies generally had low mite loads (see Supplementary Data 1 for mite levels).

**Figure 4.**
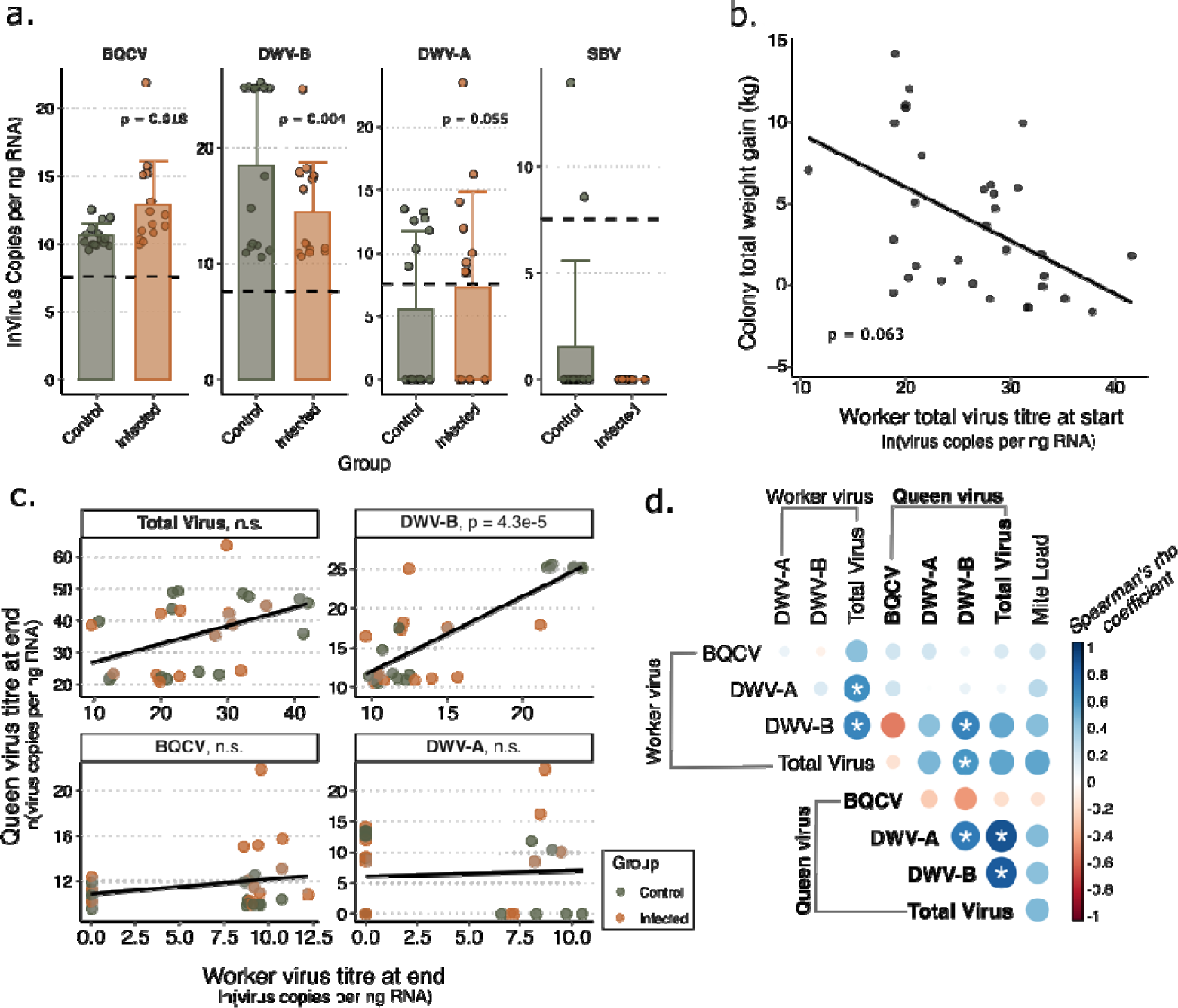
Virus titres in field experiment queens and workers. Control queens (*n* = 15) were injected with saline and infected queens (*n* = 14) were injected with a combination of BQCV and DWV-B. Queens were kept in nucleus colonies at two field sites in the lower mainland of British Columbia and sacrificed 7 weeks after injection. See Table 2 for detailed statistical reporting. **a.** Bars show mean values and error bars represent standard deviation. In the infected group, BQCV and DWV-A titres increased while titres of DWV-B decreased. There were no differences in the titres of SBV between the two groups. The dashed horizontal lines represent the RT-qPCR quantificatio threshold. **b.** The total weight gained by a colony over the 7 weeks was associated with the levels of virus present in the workers at the start of the experiment. **c.** The level of DWV-B infection in the queens was strongly correlated with the level of DWV-B present in the workers of the colony at the end of the experiment, and the trend appears to be driven by control group colonies. **d.** Correlation matrix showing the Spearman correlation between worker virus titres, queen virus titres, and colony mite counts. Asterisks represent relationships that have p < lil/n (Bonferroni correction for multiple hypothesis testing).

**Table 2.**
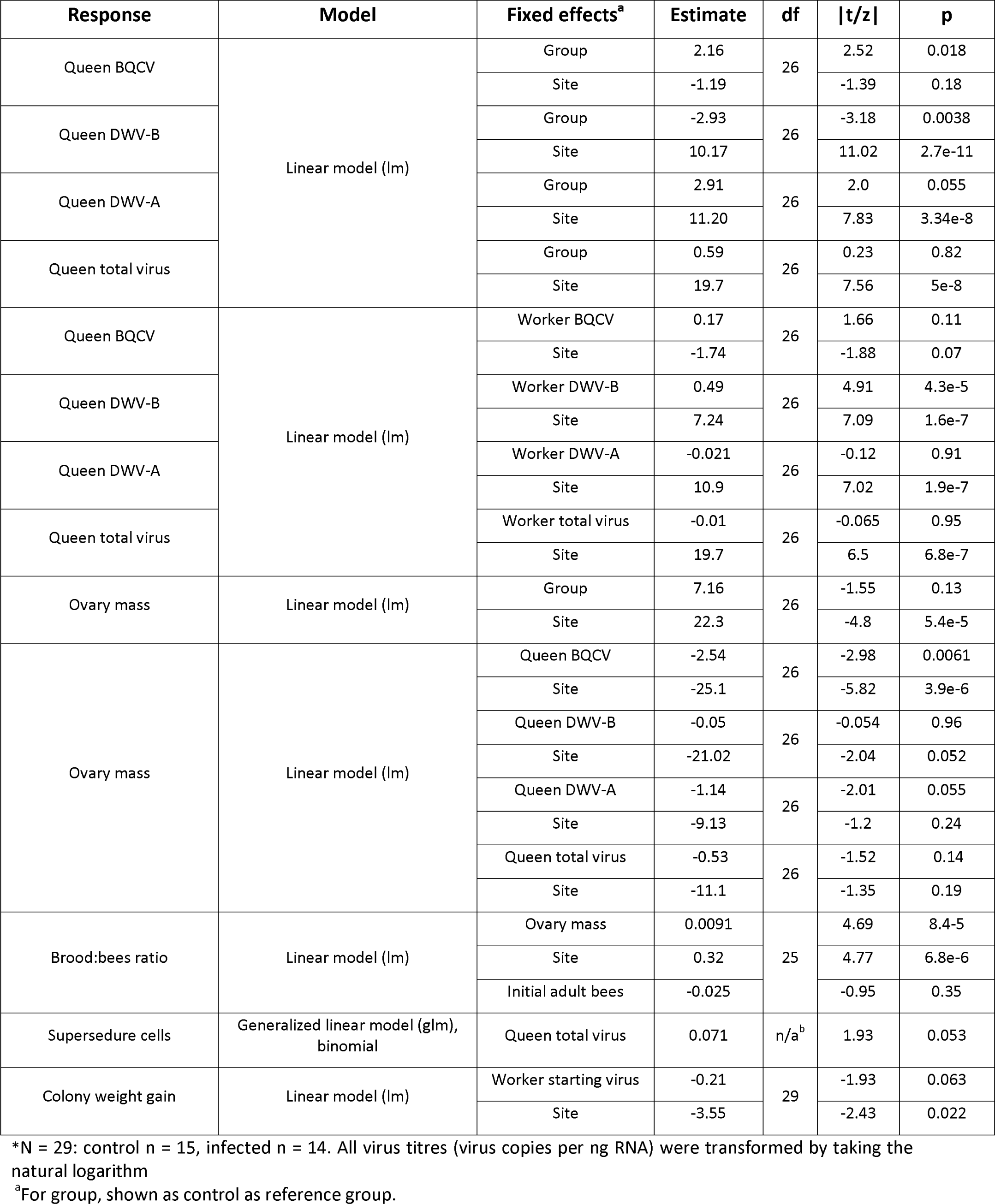
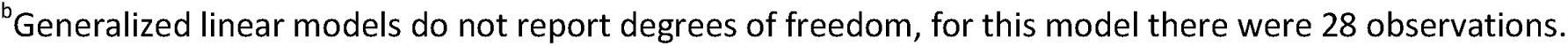
Statistical parameters for field experiment*.

| Response | Model | Fixed effects<sup>a</sup> | Estimate | df | \|t/z\| | p |
| --- | --- | --- | --- | --- | --- | --- |
| Queen BQCV | Linear model (lm) | Group | 2.16 | 26 | 2.52 | 0.018 |
|  |  | Site | -1.19 |  | -1.39 | 0.18 |
| Queen DWV-B |  | Group | -2.93 | 26 | -3.18 | 0.0038 |
|  |  | Site | 10.17 |  | 11.02 | 2.7e-11 |
| Queen DWV-A |  | Group | 2.91 | 26 | 2.0 | 0.055 |
|  |  | Site | 11.20 |  | 7.83 | 3.34e-8 |
| Queen total virus |  | Group | 0.59 | 26 | 0.23 | 0.82 |
|  |  | Site | 19.7 |  | 7.56 | 5e-8 |
| Queen BQCV | Linear model (lm) | Worker BQCV | 0.17 | 26 | 1.66 | 0.11 |
|  |  | Site | -1.74 |  | -1.88 | 0.07 |
| Queen DWV-B |  | Worker DWV-B | 0.49 | 26 | 4.91 | 4.3e-5 |
|  |  | Site | 7.24 |  | 7.09 | 1.6e-7 |
| Queen DWV-A |  | Worker DWV-A | -0.021 | 26 | -0.12 | 0.91 |
|  |  | Site | 10.9 |  | 7.02 | 1.9e-7 |
| Queen total virus |  | Worker total virus | -0.01 | 26 | -0.065 | 0.95 |
|  |  | Site | 19.7 |  | 6.5 | 6.8e-7 |
| Ovary mass | Linear model (lm) | Group | 7.16 | 26 | -1.55 | 0.13 |
|  |  | Site | 22.3 |  | -4.8 | 5.4e-5 |
| Ovary mass | Linear model (lm) | Queen BQCV | -2.54 | 26 | -2.98 | 0.0061 |
|  |  | Site | -25.1 |  | -5.82 | 3.9e-6 |
|  |  | Queen DWV-B | -0.05 | 26 | -0.054 | 0.96 |
|  |  | Site | -21.02 |  | -2.04 | 0.052 |
|  |  | Queen DWV-A | -1.14 | 26 | -2.01 | 0.055 |
|  |  | Site | -9.13 |  | -1.2 | 0.24 |
|  |  | Queen total virus | -0.53 | 26 | -1.52 | 0.14 |
|  |  | Site | -11.1 |  | -1.35 | 0.19 |
| Brood:bees ratio | Linear model (lm) | Ovary mass | 0.0091 | 25 | 4.69 | 8.4-5 |
|  |  | Site | 0.32 |  | 4.77 | 6.8e-6 |
|  |  | Initial adult bees | -0.025 |  | -0.95 | 0.35 |
| Supersedure cells | Generalized linear model (glm), binomial | Queen total virus | 0.071 | n/a<sup>b</sup> | 1.93 | 0.053 |
| Colony weight gain | Linear model (lm) | Worker starting virus | -0.21 | 29 | -1.93 | 0.063 |
|  |  | Site | -3.55 |  | -2.43 | 0.022 |

\*N = 29: control n = 15, infected n = 14. All virus titres (virus copies per ng RNA) were transformed by taking the natural logarithm
<sup>a</sup>For group, shown as control as reference group.
<sup>b</sup>Generalized linear models do not report degrees of freedom, for this model there were 28 observations.

### Field experiment – supersedure cells, ovary mass, and brood area

There was no difference in the ovary masses of the queens between the two groups, which was unsurprising given the similar levels of infection, but as we observed in the cage trial, there was a negative correlation between ovary mass and the levels of DWV-A and BQCV after controlling for the effect of site (Figure 5a, see Table 2 for statistical details). To investigate whether ovary mass, as a proxy-metric for queen fertility, could be directly linked to colony population, we assessed the total brood and adult bee area present in the colonies over the course of the experiment. We found that at the termination of the experiment (the time point closest to when queens were sampled), the ratio of brood to adult bees was positively associated with ovary mass (Figure 5b). This relationship appears to be driven by the correlation between ovary mass and increasing brood area, as there is not a significant relationship between ovary mass and adult bee population after controlling for the number of adult bees present at the start of the experiment (see Supplementary Figure 2). Premature supersedure is a common symptom associated with beekeeper-identified poor or failing queens, and we found that colonies with more heavily infected queens were indeed more likely to build at least one supersedure cell during the experiment, though the statistical significance of this effect was marginal (Figure 5c; binomial logistic regression/GLM, est. = 0.070, z = 1.93, p = 0.053).

**Figure 5.**
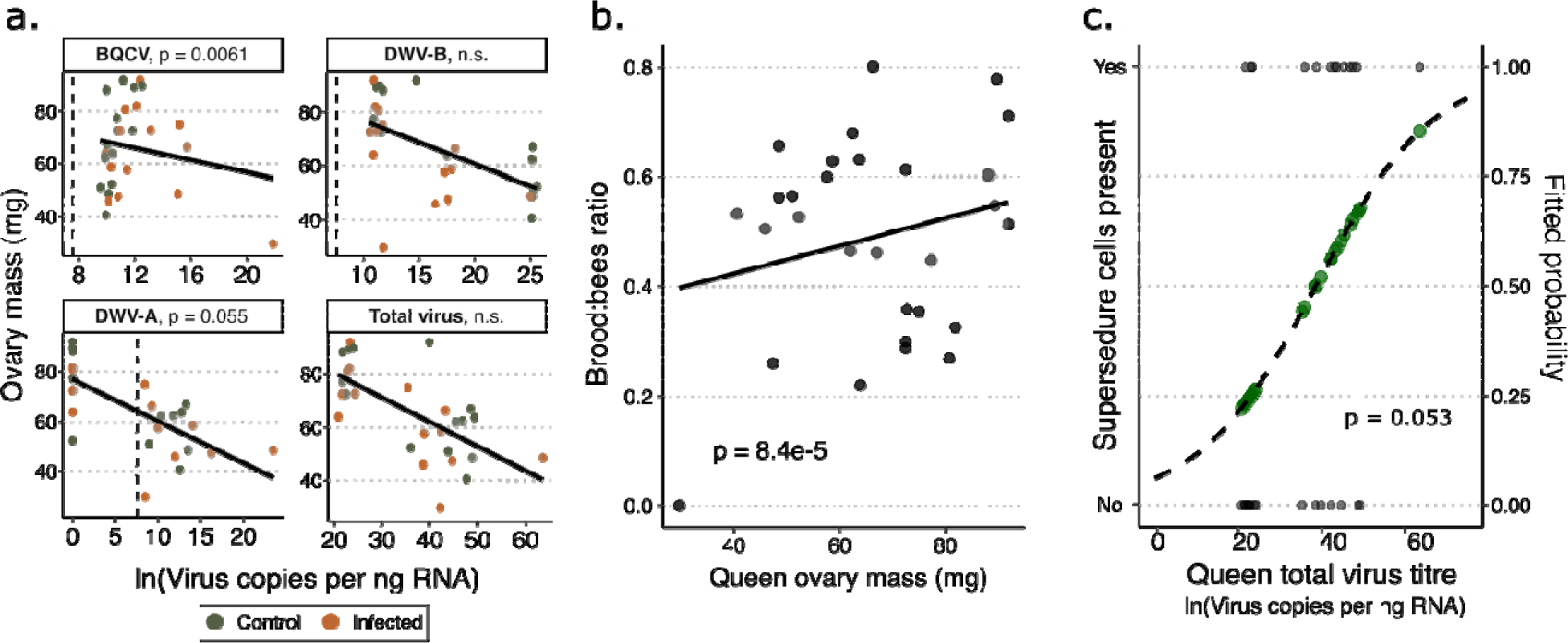
Relationships between ovary mass, virus titre, brood, and supersedure cells in field experiment queens. See Table 2 for complete statistical details. **a.** Queen ovary mass is negatively associated with levels of DWV-B, DWV-A, and total virus load. The dashed vertical lines represent the RT-qPCR quantification threshold. **b.** The ratio of brood to adult bees, measured as number of frame equivalents, present in the colony is positively associated with the ovary mass of the queens after controlling for the effect of the site and the number of adult bees present at the start of the experiment. **c.** The probability of supersedure cells being observed at some point during th experiment increases with increasing levels of virus in the queens. The green points along the curve represent the fitted values of the binomial generalized linear model.

### Field experiment – hemolymph proteome changes

While the experimental infection in the field queens did not yield significantly different viral titres between the two groups, we were nonetheless interested in investigating what proteomic changes correspond to actual virus infection level and ovary size. Quantitative proteomics analysis of hemolymph showed that, of the 1,408 protein groups quantified (1% FDR) in at least 75% of samples, 28 proteins were significantly associated with ovary mass (Figure 6, 5% FDR, Benjamini-Hochberg correction) several of which overlap with proteins differentially expressed among the groups in the cage experiment. These overlapping proteins cluster together (Euclidean distance) and are immune-related. Only four proteins were significantly associated with total viral load: an uncharacterized protein (A0A7M7TFJ8), DWV genome polyprotein (G3F401), and the *V. apis* channel protein (T0LDL0) were upregulated and phospholipase A1 (A0A7M7IQ52) was downregulated. When both ovary mass and viral load are included in the same model, though, only one protein (beta glucuronidase, A0A7M7MM50, upregulated) is significantly associated with ovary mass and none with virus, reflecting the high level of autocorrelation between the two variables.

**Figure 6.**
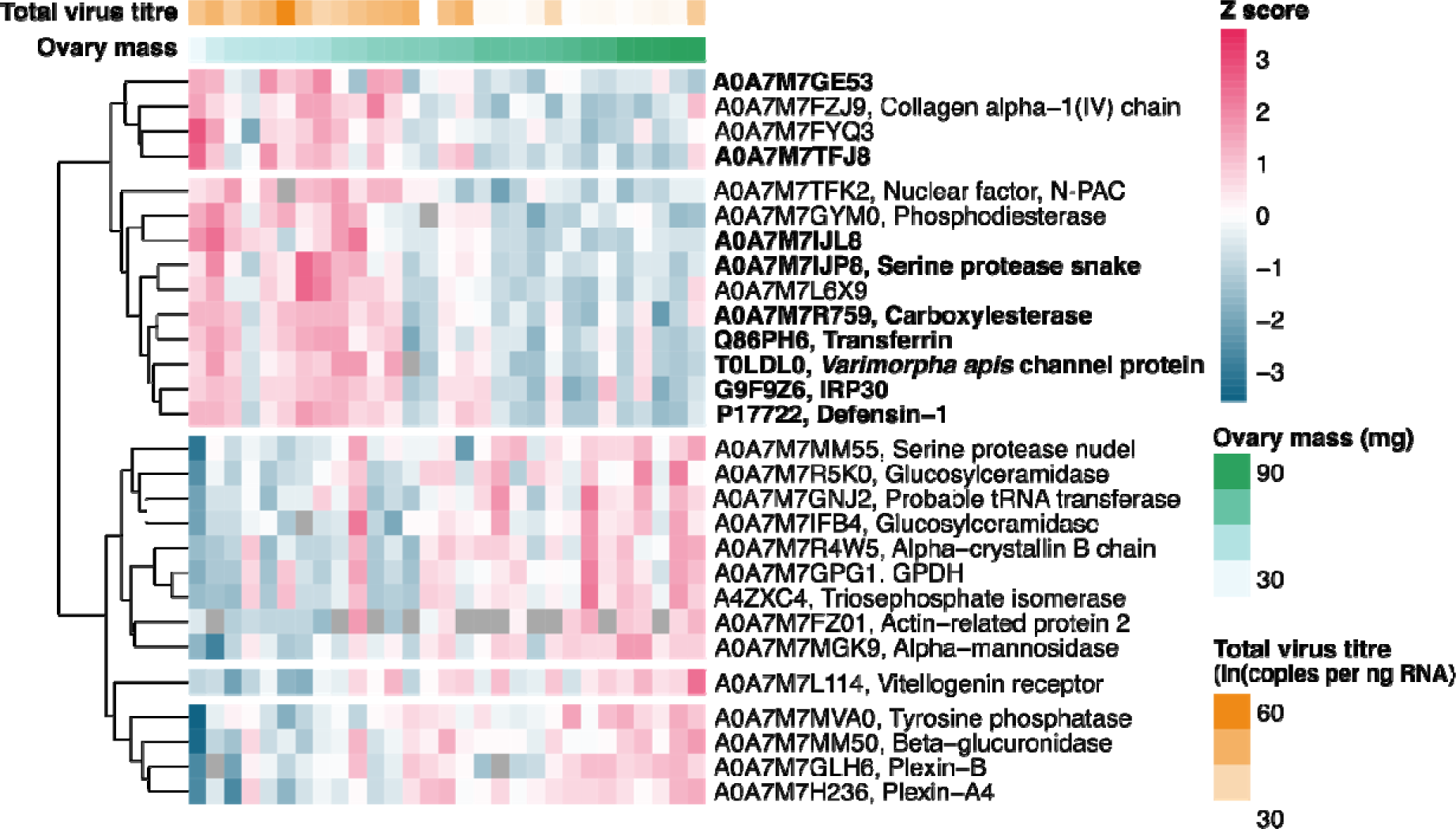
Proteins associated with ovary mass in the hemolymph of the field experiment queens. Each row represents a protein significantly associated with ovary mass (5% FDR, Benjamini Hochberg method), showing th Uniprot accession number, and each column represents a sample. Total virus titre was not included in this limma model but is shown here to illustrate the correlation between the two variables. Proteins that were not quantified in a sample are shown as grey tiles. Proteins in bold were also differentially expressed between groups in the hemolymph of the cage experiment queens (5% FDR, Benjamini-Hochberg correction). Proteins are clustered via Euclidean distance.

## Discussion

In this study, we have further substantiated the relationship between viral infection and the symptoms associated with beekeeper-identified poor queens while also investigating a potential reproductive-immunity compromise using two parallel infection experiments. In the cage experiment, we found that queens injected with infectious virus had significantly smaller ovaries and were significantly less likely to begin laying eggs again after infection, while queens that were injected with inactivated virus had marginally smaller ovaries and had no difference in their likelihood to recommence laying. However, both experimental groups exhibited a similar immune response, as determined by upregulation of immune proteins in the hemolymph. And, across all three groups, higher levels of virus were associated with smaller ovaries and fewer eggs laid. The field experiment became a correlational rather than manipulative study, as our experimental infection surprisingly did not alter the overall virus load of the queens (as measured at the end of the experiment). However, the total virus load of the queens, regardless of the group, was indeed negatively associated with ovary mass and positively associated with the presence of supersedure cells, suggesting that there are real impacts of viral infections on queen productivity and possibly worker perception of the queen. This work connects the proxy-metric for fertility of ovary mass to effects observable at the level of reproductive output (egg-laying) and at the colony level (relative amount of brood). This validates using ovary mass as a metric for current reproductive health, if compared among similar queens in the same environment. Importantly, though, the ovary masses of queens of different ages, sampled at different times, from different locations, and especially those that have been caged and/or transported for different lengths of time, should not be directly compared. In a previous study, we found that the small ovary mass of queens which had been caged for shipping rebounded after being placed back into colonies (McAfee et al., 2021).

We were surprised to find that the queens we attempted to infect in the field trial did not actually have higher viral titers than the control group. This contrasts with the cage experiment, where infected queens did have elevated viral loads using the same inoculum; therefore, we know our technique is not to blame. This could be due to the lengthened time frame, which could have allowed queens to either recover from the acute effects of infection or acquire new infections from contact with workers. However, it is likely that queens in the cage trials were also nutritionally stressed to some degree compared to queens in the field trial. This could have exacerbated the effects of the experimental infection given that the egg laying output of queens has been shown to be directly related to the nutrition available to her attendants (Fine et al., 2018). Moreover, oogenesis is more severely disrupted in queens kept in cages outside of a colony compared to queens caged inside of a colony, likely due to improved nutrition provided to queens in the colony setting (Aamidor et al., 2022). Cages had fewer (∼100) workers present to act as attendants and were supplied with pollen patty supplement, which is less nutritious than the bee bread present in the colonies (Watkins de Jong et al., 2019). Moreover, the attendants that were present in the cages aged beyond the optimal age of nurse bees during the experiment (Lass & Crailsheim, 1996). While the nutritional stress queens experience in cages is somewhat artificial, queens could in theory also become nutritionally stressed in colonies during periods of dearth, if worker populations have dwindled due to disease or other stressors, or if mismanagement leads to suboptimal age structure for brood and queen care. We therefore argue that our findings from the cage trial still have the practical utility of highlighting the range of effects that are theoretically possible to occur in colonies.

We endeavored to further investigate whether the impact of virus exposure on reproductive output was due entirely to the pathogenic effects of viral infection or if it was the result of a reproduction immunity trade-off, which assumes resources are limited to some extent. Such a trade-off is therefore expected to be exacerbated by the reduced nutrition present in the cage environment. Simply comparing the reproductive output of queens challenged with live pathogens fails to fully address this question, given that it is impossible to decouple the effects of the pathogen itself from the effects of the immune response that it has induced. However, by injecting queens with inactivated virus, we were able to isolate immune effects from pathogenicity. The moderated reduction in ovary size in the queens receiving the inactivated inoculum suggests that honey bees have partially, but not totally, escaped the reproduction-immunocompetence trade-off. This partial escape is likely due in part to social insect queens’ access to far more, but not infinitely more, nutritional resources than solitary reproductive individuals. Additionally, the presence of multi-functional proteins that are involved in both reproduction and immunity (e.g., IRP30; see below) are a tantalizing potential mechanism for reducing the overall cost to sustaining both tasks simultaneously, since resources become effectively shared via common pathways.

Honey bees exhibit a surprising scarcity of immune related genes, possessing only about one-third of those found in *Drosophila* (Evans et al., 2006). However, Drosophila is also far better-studied and there could be unrecognized immune proteins which are either uniquely multi-functional or are yet uncharacterized in social insects. One such protein is IRP30 (immune response protein 30), which was the top upregulated protein in both the live virus and inactivated groups in the cage experiment and negatively associated with ovary mass in hemolymph of the field experiment queens. IRP30 is characterized as an immune effector unique to Hymenoptera and is upregulated in response to a variety of immune challenges including bacteria (Randolt et al., 2008), virus (Fujiyuki et al., 2009), and double-stranded (ds)RNA (Nunes et al., 2013). More recently, it has robustly been shown to also be tied to reproduction—in both workers and queens in bumble bees and honey bees, the expression of IRP30 increases between 20-to 90-fold in egg-laying individuals (Dong et al., 2020). Vitellogenin is another hallmark multi-functional protein, which is not only the major egg yolk protein precursor responsible for provisioning nutrients to developing oocytes, but also both a well-established player in innate immunity in a wide variety of organisms. It functions as a pathogen-associated molecular pattern (PAMP) recognition protein (Leipart et al., 2022; C. Sun & Zhang, 2015; W. Sun et al., 2020), acts as a carrier of pathogen fragments in transgenerational priming (Harwood et al., 2019), and possesses antioxidant functions (Corona et al., 2007; Seehuus et al., 2006). Vitellogenin has also been shown to be upregulated in response to *Varimorpha ceranae* infection, perhaps contributing to a compensatory increase in antioxidant capacity (Alaux et al., 2011). The dual role of these proteins in both reproduction and immunity means that a resource allocation conflict should be mitigated, since resources invested in producing these proteins can support both processes.

If the intermediate phenotype of queens receiving inactivated virus was caused by an immune response to the viral particles alone, we would expect queens receiving the live virus to have a stronger response since the number of live viral particles proliferated far beyond the number contained in the single dose of inactivated virus. This is the response we observed for ovary masses, but not immune proteins. The patterns of immune protein expression indicate that both treatments appear to stimulate a very similar immune response, both in the identity of the effectors and the magnitude of expression. Even though this was a viral infection experiment, all the immune-related proteins we identified as significant are canonically associated with the immune response to bacterial and fungal infection. This is unsurprising, given that it has been repeatedly shown there is significant cross-talk among the various immune response pathways, and so-called “antimicrobial” effectors are robustly upregulated in response to viral infection in honey bees (Durand et al., 2023; McMenamin et al., 2018). Interestingly, while proteins were differentially expressed between groups in the cage experiment, there were no proteins significantly associated with the total viral load. While this is likely due at least in part to easier statistical ability to detect differences between discrete groups rather than among variable, continuous data with somewhat limited sample numbers, it could also indicate that a perturbation from an individual’s background virus infection in an acute time frame is more impactful to the immune system and ovary mass than absolute virus load (which includes some level of pre-existing infections). Several of these proteins were associated with ovary mass in the field experiment, highlighting how interconnected ovary mass and response to viral infection is. It is not clear why the immune response in the queens receiving live virus did not have a larger magnitude than the inactivated group, but it is possible that there is an upper limit to immune inducibility which we surpassed with our treatments.

The *V. apis* protein that was identified in both experiments is intriguing. *V. apis* has been known to frequently co-infect both colonies and individuals with BQCV, and has been shown to act synergistically to increase mortality of adults bees beyond infection with either pathogen individually (Bailey et al., 1983), although the way in which these pathogens interact is still unknown. This protein was the only one identified in its protein group, and several other proteins from this pathogen were quantified (although not differentially abundant) which suggests that this is a legitimate finding. The *V. ceranae* proteome was included in the search database, indicating these are not mis-identified microsporidia proteins either. It is unlikely that the viral inoculum was contaminated with *V. apis* spores after ultra-centrifugation, given the large size difference between virus particles and spores. Because we also see this protein in the inactivated group, this suggests that perhaps there is something about stimulating the immune system, either through natural BQCV infection or another stimulant like inactivated virus, which may actually help *V. apis* invade and proliferate. More research about the relationship between these pathogens and immune stimulation is needed to address this possibility.

Queen “failure” in the field has previously been associated with higher levels of viruses, and we were able to recapitulate symptoms of failure with experimental infections; however, the results of our study also suggest that queen health is intrinsically tied to the circumstances of her colony environment. For instance, levels of virus infection in the colony and in the queen likely have a two-way interaction. Here, we show that in the colony context, it appears that the levels of DWV-B in the workers have a stronger influence on the queen’s DWV-B titre than actual injection with the virus, despite the layers of social immunity (Evans et al., 2006) that should conceivably shield her from exposure. The presence of *Varroa* in the colony adds an additional influencing factor, given that DWV is transmitted by mites (de Miranda & Genersch, 2010) and we found that levels of DWV and mite load are positively correlated, although not significantly. Given that the colonies with the highest levels of DWV-B in the workers and queens were control colonies, it seems unlikely that the relationship between queen and worker infections is driven by infected queens increasing the titres in their colonies (though it has previously been shown that virus infection can be vertically transmitted from queen to progeny; Chen et al., 2006; de Miranda & Fries, 2008; Ravoet et al., 2015). Many of the common queen failure symptoms can actually be attributed to factors external to the queen, as illustrated by Lee et al. (2019), who found that switching queens with poor or “failing” brood patterns into colonies with “good” brood patterns significantly improved the brood pattern of the “bad” queen.

Therefore, while viral infection is clearly tied to queen reproductive output, it seems likely that adequate care from her colony can at least partially ameliorate these effects. This is consistent with our hypothesis that honey bee queens may have an increased threshold beyond which a trade-off manifests, rather than a simple presence or lack of a reproduction-immunity trade-off. This could be mediated by multiple factors which are not mutually exclusive: (1) a decreased cost to activating both processes due to the presence of Hymenopteran-specific, multi-functional proteins which function in both reproduction and immunity; (2) the rewiring of specific hormones, which act as a discrete switch between the two processes; or (3) access to an unusually high amount of energetic resources among invertebrates due to social provisioning. Given that the latter element is highly dependent on the health status of the colony, and we found that a compromise between reproduction and immune activation may be partially responsible for some of these effects, this represents a further way in which what we observe as symptoms associated with a queen could be tied to the conditions of the colony. Aside from several issues with queen reproductive health that can be directly attributed to poor mating, aging, or reduced sperm quality, laying blame solely on the queen in cases of queen “failure” belies a significant part of the story involving the health, infection, and nutrition status of the colony.

## Conclusion

Our results indicate that the effects of viral infections in queens translate to the real-world symptoms about which beekeepers are concerned, and these effects are partially, but not entirely, governed by a reproduction-immunity trade-off. Attempted or premature supersedure is an often-cited indicator of queen failure by beekeepers, which we have now shown to be directly correlated to queen virus infection. The acute effects of experimental virus infection were magnified in our laboratory trials, where nutritional stress was likely an exacerbating factor, compared to the field trial where colonies with excess food availability could provide queens with more adequate nutrition. Forage dearth and therefore nutritional stress can, however, also occur in the field, and our data suggest that effects of virus infection on queens may be worsened under such conditions. These data collectively underscore the likelihood that a reproductive-immunity trade-off may be at play, though possibly moderated by the vast amount of nutritional resources available to social insect queens. This work provides further evidence that virus infections are directly related to symptoms beekeepers identify in poor queens in the field, highlights the ways in which queen health and her colony environment are inextricably related, and underscores the importance of managing virus levels in beekeeping operations.

## Methods

### Cage experiment

Young, mated queens (reared from closely related mothers) that had commenced laying within the previous week were couriered overnight in JZBZ battery boxes filled with nurses from a British Columbia queen producer (Wild Antho). Queens were received in three batches, each 2 weeks apart, starting in the middle of July. Queens were placed in queen monitoring cages (QMCs) (Fine et al., 2018) along with approximately 11 grams of worker bees (∼100) collected from a brood frame of a single colony maintained at the University of British Columbia (UBC). The cages were kept in an incubator at 34 °C, with a large water pan to increase humidity, and were supplied *ad libitum* with fondant (Ambrosia), protein patty (Global Patties), and tap drinking water. A total of *n* = 9 “baseline” queens, three from each of the three batches received, were anesthetized and frozen immediately upon receipt. Each QMC contained two “egg laying plates” that are made up of 264 hexagonal cells in the dimensions of natural brood comb (Fine et al., 2018). These plates were cleaned and/or replaced every 48-72 hours to provide queens with continuous space to lay. Queens were allowed to acclimate to the cages and confirmed to be consistently laying for 6 days before injection. Queens were then injected on Day 6 and observed for 7 more days before sampling. For each batch, queens were randomly assigned to each experimental group (saline injected control, live virus infected, and UV-inactivated virus injected) such that the experimenter (Chapman) was blind to group identifiers. In total, *n* = 9 queens were tested in each group, spread equally across the three experimental batches (3 in each batch). Blinding was maintained until all data had been collected.

### Field experiment

Young, mated queens (*n* = 32) were obtained as a single batch of sister queens from the same producer as for the cage experiment and introduced into small, five-frame colonies (referred to by beekeepers as nucleus colonies or “nucs”). The nucs were established on June 7, 2023 almost exclusively from a commercial source of bees (packages obtained from Kintail and Arataki suppliers in New Zealand) that were previously hived in March and allowed to grow until June. Stock imported from New Zealand into Canada is required to have less than 1% *Varroa* mites (mites per 100 bees), but an oxalic acid (3.5% in 50% syrup) mite treatment was applied at hiving as a precautionary measure. Some additional stock was pulled from overwintered colonies maintained at the University of British Columbia that had sufficiently low (<1%) mite loads and an absence of brood disease symptoms. Nucs were produced such that each colony had one frame of honey, two frames of brood, one frame with space for laying (drawn, irradiated comb), and one frame of plastic foundation. Adult bee populations were equalized as much as possible during nuc production but they were further equalized 3 days after queen introduction by adding young adult bees to smaller colonies. The experiment was conducted at two sites 2 km apart (16 nucs at each site) such that no nuc remained at the same site as its parent colony. Nucs were provided with protein supplement (Global Patty) and fondant (Hive Alive) and replenished weekly throughout the experiment to minimize potential nutritional differences among colonies. On July 7, European foulbrood disease was detected in a subset of colonies, therefore all colonies at both sites were treated with Oxytet-25 (oxytetracycline antibiotic) according to the label rate to minimize impact of the disease on the experiment while still managing all colonies equally.

Two weeks after the nucs were produced, baseline colony weights were recorded (supplemental feed was removed prior to weighing), a sample of 15 randomly selected workers from a brood frame was taken for initial worker viral analysis, and both sides of each frame were photographed for subsequent brood area and adult bee coverage calculations. The same day, queens were retrieved then randomly and blindly assigned (with the help of a colleague not otherwise involved in the experiment) to either the saline injected control or live virus infected group, with balanced groups within each site. Once injected, the queens were returned to their respective colonies within 1-2 hours. From that point on, colony weights were recorded weekly and frames were photographed weekly or biweekly (June 22, June 29, July 6, July 19, and Aug 2). During photographing, incidence of brood disease (number of affected cells) was noted and the presence or absence of supersedure cells was recorded before the cells were destroyed to prevent queen loss. During the course of the experiment, 11 colonies grew sufficiently to require an additional box of frames (consisting of a mixture of foundation and irradiated drawn comb), which were weighed and subtracted from the subsequently measured colony weights. At the end of the experiment (Aug 10), we sampled the experimental queens, 15 randomly selected workers from a brood frame (for final worker viral analysis), and ½ cup of bees (∼300 adults) for *Varroa* mite counts using alcohol wash method (Jong et al., 1982). A total of 29 queens persisted to the end of the experiment, as in the final 2 weeks one queen was lost by accident and two were superseded. The operators (Chapman and McAfee) remained blind to experimental groups until all data were collected, including image analysis.

Photographs of frames were analyzed using ImageJ to calculate the total area of adult bee coverage and brood coverage. In cases where adult bee coverage was too dense to adequately assess the brood, each side of the frame was photographed, then the bees were gently agitated to expose the brood for a second set of photographs. Top bars were labelled with the colony ID and frame number to associate photos with colonies. In ImageJ, bee area and brood area were traced and calculated, then expressed as a fraction of total frame area, which were summed for each colony.

### Virus isolation and propagation

Samples of homogenized honey bees from individual colonies that had been stored at −80°C from previous projects were tested by RT-qPCR to identify samples with the highest concentration of the target virus. Those with the highest virus purity relative to other commonly detected viruses in our samples (DWV-B, DWV-A, IAPV, BQCV and SBV) were used for the primary extracts as described by de Miranda et al., 2013. These extracts were passed through a Corning syringe filter (0.45 µm) to remove bacteria. Primary extracts were used to make 100 µL serial dilutions; DWV primary extracts were diluted with 90 µL of 0.5M potassium phosphate buffer solution (pH 7.5), BQCV primary extracts were diluted with 0.01 M potassium phosphate buffer (pH 7.0), and three concentrations (10^5^, 10^4^, and 10^3^ gene copies per µL) were used to inoculate pupae.

Pupae that contained low background virus levels were obtained from honey bees that had been initiated from Australian package bees (Aussie bees® sourced from Tasmania) hived on new plastic foundation and held in isolated apiaries in Canada (Star lake Manitoba 49°45’08“N 95°15’24“W and Red Deer Lake Ontario 50°00’47“N 94°11’26“W) in areas where no other managed honey bees occur. Colonies were sampled using sticky boards and alcohol washes to confirm the absence of *Varroa* mites. Honey bee pupae collected at the white-eyed to pink-eyed stage were then used to propagate the viruses. A Hamilton® syringe with a 30GxK gauge sterile needle was mounted on a Schley Compact MDL II® insemination stand and then the target virus solution was inoculated into the pupae through the lateral side of the abdomen between the 2^nd^ and 3^rd^ tergites. One µL of virus solution was injected per bee. Inoculated pupae were then placed in a covered Petri dish with paper towel dampened with distilled water, with 20 pupae per dish and incubated at 31.5°C for 5 days. Pupae were checked regularly, and distilled water was added as required with any dead pupae removed from the dishes. Any pupae that reached the adult stage prior to the end of 5 days were stored at −80°C for later processing. After incubation, all healthy pupae (or adults) from each dose (10^5^, 10^4^, or 10^3^ virus particles) were collected in three 50 mL VWR centrifuge tubes. The pupae were stored at −80°C until further processing or homogenized immediately after collection.

### Virus Purification

Extraction buffer, which varied with type of virus, was first added to the propagated virus samples. DWV extraction buffer consisted of 0.5 M potassium phosphate buffer (pH 7.5) and 0.2% ascorbic acid. BQCV buffer consisted of 0.01 M potassium phosphate buffer (pH 7.0) and 0.02% ascorbic acid. To make primary extracts, 2 mL of extraction buffer and 0.5 mL chloroform were added per 1 g of homogenate and samples were kept on ice in a fume hood. Each sample was vortexed for 30 seconds, then centrifuged at 8,000 g (8,700 rpm, Eppendorf Centrifuge 5430, placed in the fridge) for 10 min at 15°C. From each tube, the supernatant was collected into a 5 mL centrifuge tube based on sample origin. Samples were stored at −80°C until further purification.

High speed centrifugation was done to further purify the virus samples. The primary extracts were thawed and added to 35 mL Thermo Scientific™ PA Ultracrimp Tubes, with the rest of the volume filled with more extraction buffer. The tubes were centrifuged at 15 °C for 3 hours, at 75,000 g (27,200 rpm, Sorval Discovery 100SE, Sorval T-865 Rotor). The supernatants were discarded, and the pellets were placed in new 5 mL centrifuge tubes. DWV was resuspended in 5 mL of 0.5 M potassium phosphate buffer (pH 7.5), and BQCV in 0.01 M potassium phosphate (pH 7.0). The solutions were mixed using a spatula and via vortex. The samples were left overnight at 4°C and then mixed gently in the morning.

The samples were split into 2 mL microcentrifuge tubes to be centrifuged for 15 min at 4°C, at 8,000 g (8,700 rpm, Eppendorf Centrifuge 5430, placed in the fridge). The supernatants were collected into a new 5 mL centrifuge tube based on sample. The samples were then passed through a 0.45 µm syringe filter into new 5 mL centrifuge tubes, and 20 µL of each sample was taken for viral analysis to determine the final concentrations and purities.

The final extracts had the following purities: DWV-B was 99.99% pure (with other components consisting of 0.000006% DWV-A, 0.00% IAPV, 0.00% BQCV, and 0.000006% SBV), and BQCV was 99.88% pure (with other components consisting of 0.02% DWV-B, 0.02% DWV-A, 0.05% IAPV and 0.02% SBV).

### Injection of queens

Queens were anesthetized with carbon dioxide for 3 minutes (starting when the queen ceased all movement) before being injected on the dorsal side between the 2^nd^ and 3^rd^ abdominal tergites with ∼100 nL containing either sterile phosphate-buffered saline, a combination of infectious BQCV (∼1 × 10^8^ copies) and DWV-B (∼2 × 10^6^ copies), or the same combination and quantity of virus which had been UV-inactivated (cage experiment only). These doses are in line with other experimental infections in honey bees in the literature (Al Naggar & Paxton, 2020; Dubois et al., 2020; Tehel et al., 2020). Injections were performed using the FemoJet Microinjector (Eppendorf) with pulled-capillary glass needles. Injection volumes were calibrated by estimating the droplet volume using a hemocytometer grid under a compound microscope. Everything associated with the injection apparatus was wiped down or soaked in 70% ethanol before and after injections. The saline injected queens were injected first and the live virus group was injected last. To inactivate the virus mixture, a 50 µL aliquot of virus on parafilm (kept on ice) was exposed to 4 consecutive doses of 120,000 mJ of ultraviolet light using a Stratalinker UV Crosslinker (Stratagene) in a manner similar to previously published protocols (Mathew et al., 2018; Soh et al., 2023).

### Queen dissection

Queens were sacrificed by anesthetization with carbon dioxide for ∼5 min. To collect hemolymph for proteomics while avoiding potential contamination from the gut, a leg was removed from the thorax of each queen and a glass capillary was used to collect the exuded hemolymph. After hemolymph was collected, queens were decapitated and the ovaries were removed and weighed on an analytical balance. The thoraxes were stored for viral qPCR analysis. All samples were stored at −70°C until subsequent preparation.

### Proteomics sample preparation

Hemolymph samples (*n* = 36 for the cage experiment and *n* = 29 for the field experiment) were prepared for mass spectrometry essentially as previously described (Chapman et al., 2022). Between ∼5 and ∼15 µL of hemolymph which had been collected from the queens was directly precipitated by the addition of 500 µL of ice-cold 80% acetone and incubation overnight at −20°C. The precipitated protein was pelleted by centrifugation (6,000 *g*, 15 min, 4°C) and the supernatant was discarded. The protein pellet was washed twice with 500 μL of 80% acetone, then the pellet was allowed to air dry (∼5 min) before solubilization in 15 μL of digestion buffer (6 M urea, 2 M thiourea, 100 mM Tris, pH 8).

The resuspended protein was quantified using a Bradford assay (PIERCE) and then 20 μg of protein was reduced (0.4 μg dithiothreitol, 30 min) and alkylated (2 μg iodoacetamide, 20 min, in the dark). The samples were then digested overnight with 0.8 μg trypsin/LysC mixture (Promega) and diluted with 5 volumes (150 µL) of digestion buffer (50 mM ammonium bicarbonate) after the first 3 h. Digested peptides were acidified with 30 µL of 20% formic acid (to pH < 2.0) and desalted with high-capacity STAGE tips as previously described (Rappsilber et al., 2003). Eluted samples were dried (SpeedVac, Eppendorf), resuspended in Buffer A (0.1% formic acid, 2% acetonitrile), and then the peptide concentrations were determined using a NanoDrop (Thermo, 205 nm).

### Mass spectrometry data acquisition

LC–MS/MS data acquisition was performed exactly as previously described (McAfee et al., 2024). A total of 50 ng of digested peptides were injected onto the LC system in randomized order. The digest was separated using NanoElute UHPLC system (Bruker Daltonics) with Aurora Series Gen2 (CSI) analytical column (Ion Opticks, Parkville, Victoria, Australia) heated to 50°C and coupled to timsTOF Pro (Bruker Daltonics) operated in DIA-PASEF mode. A standard 30 min gradient was run from 2% B to 12% B over 15 min, then to 33% B from 15 to 30 min, then to 95% B over 0.5 min, and held at 95% B for 7.72 min, where buffer B consisted of 0.1% formic acid in 99.4% acetonitrile and buffer A consisted of 0.1% aqueous formic acid and 0.5 % acetonitrile in water. Before each run, the analytical column was conditioned with 4 column volumes of buffer A. The NanoElute thermostat temperature was set at 7°C. The analysis was performed at 0.3 μL/min flow rate.

The trapped ion mobility time-of-flight mass spectrometer (TimsTOF Pro; Bruker Daltonics, Germany) was set to parallel accumulation-serial fragmentation (PASEF) scan mode for data-independent acquisition scanning (100 – 1700 m/z). The capillary voltage was set to 1800 V, drying gas to 3 L/min, and drying temperature to 180 °C. The MS1 scan was followed by 17 consecutive PASEF ramps containing 22 non-overlapping 35 m/z isolation windows (Table 1), covering the 319.5 – 1089.5 m/z range. As for TIMS setting, ion mobility range (1/k_0_) was set to 0.70 – 1.35 V·s/cm^2^, 100 ms ramp time and accumulation time (100% duty cycle), and ramp rate of 9.42 Hz; this resulted in 1.91 s of total cycle time. The collision energy was ramped linearly as a function of mobility from 27 eV at 1/k_0_ = 0.7 V·s/cm^2^ to 55 eV at 1/k_0_ = 1.35 V·s/cm^2^. Mass accuracy was typically within 3 ppm and is not allowed to exceed 7 ppm. For calibration of ion mobility dimension, the ions of Agilent ESI-Low Tuning Mix ions were selected.

### Mass spectrometry data processing and analysis

MS data was searched using DIA-NN (v1.8.1) (Demichev et al., 2020) using “FASTA digest for library-free search” with “Deep-learning based spectra, RTs and IMs prediction”, allowing for 2 missed cleavages, selecting “Unrelated runs”, setting protein inference to “Protein names from FASTA”, and setting the neural-network classifier to “Double-pass mode”. “N-term M excision” and “C carbamidomethylation” were selected as modifications, and “Use isotopolgues”, “MBR”, and “No shared spectra” were all left checked for the algorithm. All other parameters were left set as default. The protein database was the UniProt proteome for *Apis mellifera* (based on the Amel_HAv3.1 genome assembly), plus common honey bee virus and pathogen proteins from UniProt downloaded on February 6, 2023. To this, a comprehensive list of potential protein contaminants was added (Frankenfield et al., 2022).

All subsequent analyses were performed in R (v 4.3.0). The identified and quantified protein groups from the DIA-NN search were filtered to remove contaminants and proteins identified in fewer than 75% of samples. Normalized LFQ intensity was then log2 transformed. Differential expression analysis was performed using the limma package in R (Ritchie et al., 2015). For the cage experiment, experimental batch was included as a blocking factor and group (all 4 groups including the baseline queens) was included as a fixed effect. Contrasts were defined with the control queens as reference. For the field experiment, three models were used: just ovary mass as a fixed effect, just total virus titre as fixed effect, or both. We checked if site should be included as a blocking factor, but the consensus correlation was negative, so it was not included. Empirical Bayes moderation of the standard errors was performed on the resulting outputs, and the false discovery rate was controlled to 5% using the Benjamini– Hochberg correction.

### Sample preparation for viral quantification

The thoraces from each queen were placed in separate 2 mL screw-cap homogenizer tubes containing 500 µL of Trizol reagent and three ceramic beads. The samples were homogenized (Precellys 24 homogenizer, Bertin Instruments) 2 times for 30 seconds at 6,500 rpm and placed on ice for 5 minutes between runs. The homogenate was then centrifuged for 30 seconds at 14,000 *g* and all 500 µL of supernatant was transferred to a new tube containing 500 µL of 100% ethanol for nucleic acid precipitation. For the worker samples, 5 thoraces from workers were pooled with 1 mL of Trizol reagent and 4 ceramic beads in three tubes for a total of 15 workers sampled from each colony. 250 µL of centrifuged homogenate from each of the 3 tubes was pooled with 500 µL of 100% ethanol for precipitation.

The rest of the RNA extraction was performed using the Direct-zol RNA Miniprep spin-column kit (Zymo Research), including the DNase I treatment, according to manufacturer instructions. The RNA was eluted in 50 µL of RNase-free water and quantified using a Qubit Fluorometer with the Qubit RNA High Sensitivity assay kit. A subset of samples were assessed for contaminants using a NanoDrop One spectrophotometer. The extracted RNA was stored at −70°C and thawed only once. RNA was diluted further in RNase-free water and 200 ng was used to produce cDNA. Reverse transcription was carried out using the High-Capacity cDNA Reverse Transcription Kit (Applied Biosystems), which utilizes random primers, according to manufacturer’s instructions. cDNA was kept after synthesis at 4°C for a maximum of 2 weeks.

### RT-qPCR for viral quantification

Quantitative real-time PCR was performed on a C1000 Touch Thermal Cycler with a CFX96 Touch Real-Time PCR Detection System (Bio-Rad) using SsoAdvanced™ Universal SYBR® Green Supermix (Bio-Rad). Reactions (10 µL total volume) were set in 96-well plates with 2.5 µL of cDNA at 2 ng/µL (for a total of 5 ng per reaction). Primers were included at a final concentration of 250 nM for both the forward and reverse primers, except for the SBV amplicon for which both primers were at a final concentration of 500 nM. A standard curve in triplicate was included on every plate using a single plasmid containing all the amplicons of interest. See Supplementary Data 2 for the sequences of the plasmid used for the standard curve and the primers. The standard curve spanned 6 orders of magnitude, from 9.6 × 10^8^ copies/µL to 960 copies/µL. The plasmid was dissolved in RNase-free water and the concentration was determined spectrophotometrically using a NanoDrop One. For every plate, the efficiency of the standard curve was between 90-110% and the R^2^ was ≥ 0.99. The qPCR thermocycling conditions were as suggested by the Supermix manufacturer: polymerase activation and initial denaturation for 30 sec at 95°C, followed by denaturation for 10 seconds at 95 °C, then extension for 20 seconds at 60°C for 40 cycles. A melt-curve analysis was performed after every assay following a 65-95°C gradient at 0.5°C increments every 2 seconds, and no unintended amplicons were ever observed. A “no transcript control” was included in triplicate on every plate and was never amplified. C_q_s were determined by the regression mode in the CFX Manager Software (v 2.1) rather than by a single threshold. Starting quantity (SQ) in copies was calculated within the software based on the standard curve. These values were then further processed in R (v 4.3.0) for statistical analysis. Samples with a C_q_ greater than the average C_q_ of the lowest point on the standard curve for that plate were removed. The SQ’s of the remaining samples were then averaged across replicates for each sample. Viral quantity is reported as copies/ng of starting RNA.

### Statistical analysis

All data analysis was performed in R (v 4.3.0). For analyzing the differences in virus titres and ovary mass between groups, analyzing the relationship between ovary mass and virus titre, and assessing the relationship between the number of eggs laid and the total virus titre of queens in the cage experiment, we used a linear mixed model (lmer in the lme4 package, Bates et al., 2015) in addition to lmerTest (Kuznetsova et al., 2017) to obtain p-values using Satterthwaite’s degrees of freedom method, with experimental batch as a random effect. To assess the differences in likelihood of queens beginning to lay eggs again after infection in the cage experiment, we used a generalized linear model with a binomial family function (glm in the base R “stats” package) and group as a factor (3 levels). For the field experiment, all models (see Table 2 for detailed model structures) except for the model examining the presence of supersedure cells were fit using a simple linear model including site as a fixed effect (2 levels). The supersedure cell model was fit with generalized linear model with a binomial family function (glm in base R “stats” package) and site as a fixed effect. All models were assessed for goodness-of-fit using the DHARMa package (Hartig, 2022) which utilizes simulation-based residuals to check for typical model misspecification problems such as over/underdispersion and residual autocorrelation.

## Supporting information

Supplemental Data 1 - metadata

Supplmental Data 2 - qPCR sequences

## Data availability

All proteomics raw data, search results, and search parameters are available on MassIVE (www.massive.ucsd.edu, accession MSV000094113 for the field experiment and MSV000094114 for the cage experiment) which are associated with Figures 3, 6 and Supplementary Figure 1. Sample metadata, including viral abundances, are available in Supplementary Data 1. The sequence of the plasmid used for the standard curve and primers used for the viral RT-qPCR assay are available in Supplementary Data 2. Source code underlying figures and data analysis are available freely upon request.

## Acknowledgements

Honey bee research in LJF’s group is supported by an NSERC Discovery Grant. Mass spectrometry infrastructure is supported by the Canada Foundation for Innovation, Genome Canada and Genome BC (264PRO, 374PRO) and computational infrastructure is supported by a Digital Research Alliance of Canada Resource Allocation to LJF. A Project *Apis* m. grant to AM, AC, LJF and DRT funded this research. We would like to acknowledge Emily Huxter (Wild Antho) for providing the queens, Scott Covey for help developing the RT-qPCR methods, and the UBC proteomics core facility team—Jason Rogalski, Renata Moravcova, and Jeanne Yuan—for running the mass spectrometry samples, instrument maintenance, and technical expertise. Mention of trade names or commercial products in this publication is solely for the purpose of providing specific information and does not imply recommendation or endorsement by the U.S. Department of Agriculture. USDA is an equal opportunity provider and employer.

## Author contributions

AM and AC conceptualized the experiments and analyses with input from LFJ and DRT. AC wrote the first draft of the manuscript, made the figures, and interpreted the data with assistance from AM. LFJ, DRT, RWC, JF, AM, ZR and KP provided editing assistance. AC conducted the cage experiment and AM conducted the field experiment with assistance from AC. JF provided the queen monitoring cages and provided guidance on their use. RWC, KP, and ZR produced the purified virus for use in experimental infections. AC conducted the proteomics and RT-qPCR virus analysis. Grants supplied to AC, AM, LJF, and DRT funded the work.

