## Supplementary material for "Common viral infections inhibit egg laying in honey bee queens and are linked to premature supersedure": Supplmental Figures

**Supplementary figures**

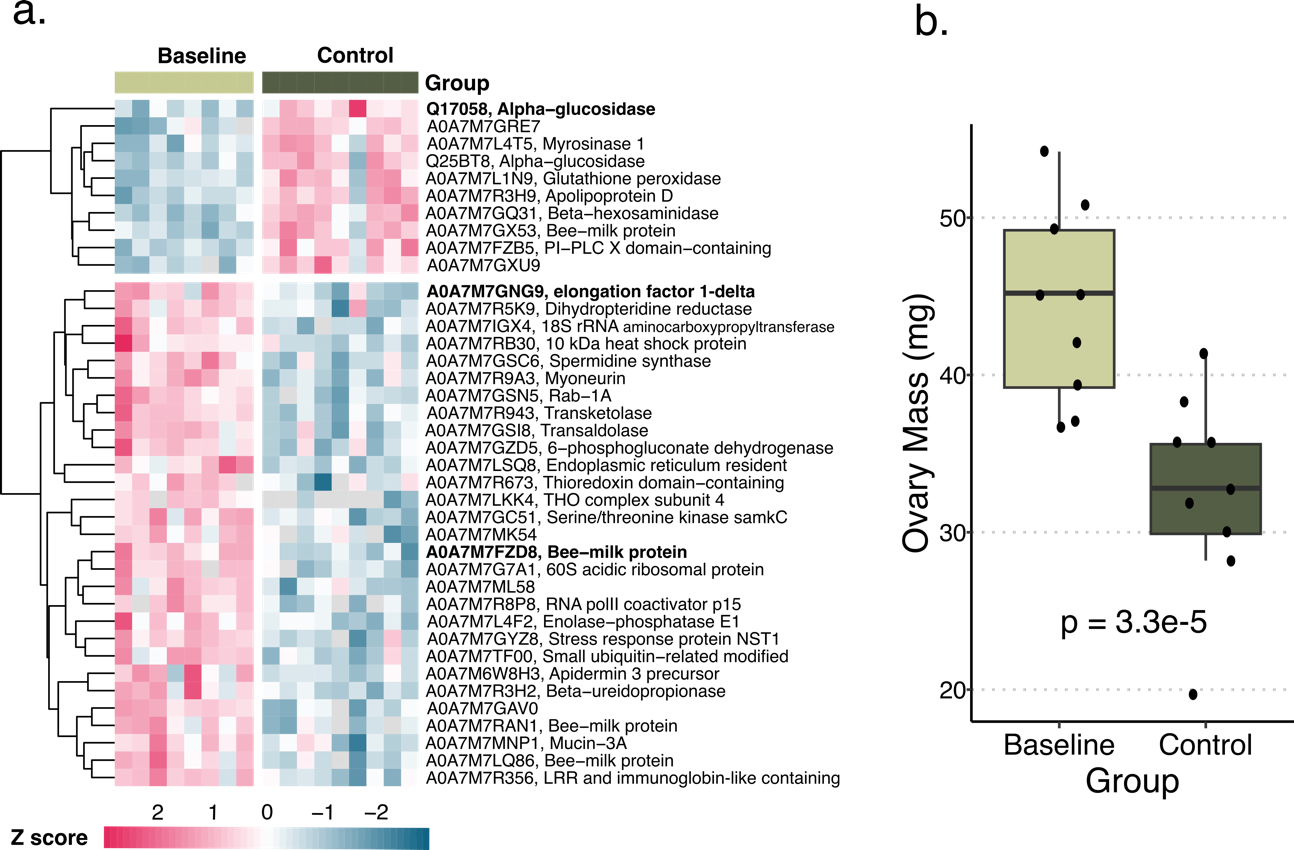

**Supplementary Figure 1. Effects of cage environment on ovary mass and protein expression.** Each row represents a protein differentially expressed between the baseline queens and the control queens, showing the Uniprot accession number, and each column represents a sample. Proteins that were not quantified in a sample are shown as grey tiles. Proteins in bold were also differentially expressed between the control and live virus group queens (5% FDR, Benjamini-Hochberg correction). Proteins are clustered via Euclidean distance. **a.** While 39 proteins are differentially expressed between the baseline and control queens, only 3 overlap with the proteins associated with the effects of the experimental injections. **b.** The ovaries of control queens are smaller than baseline queens (linear model est. = -12.86, t = -5.99, p = 3.3e-5).

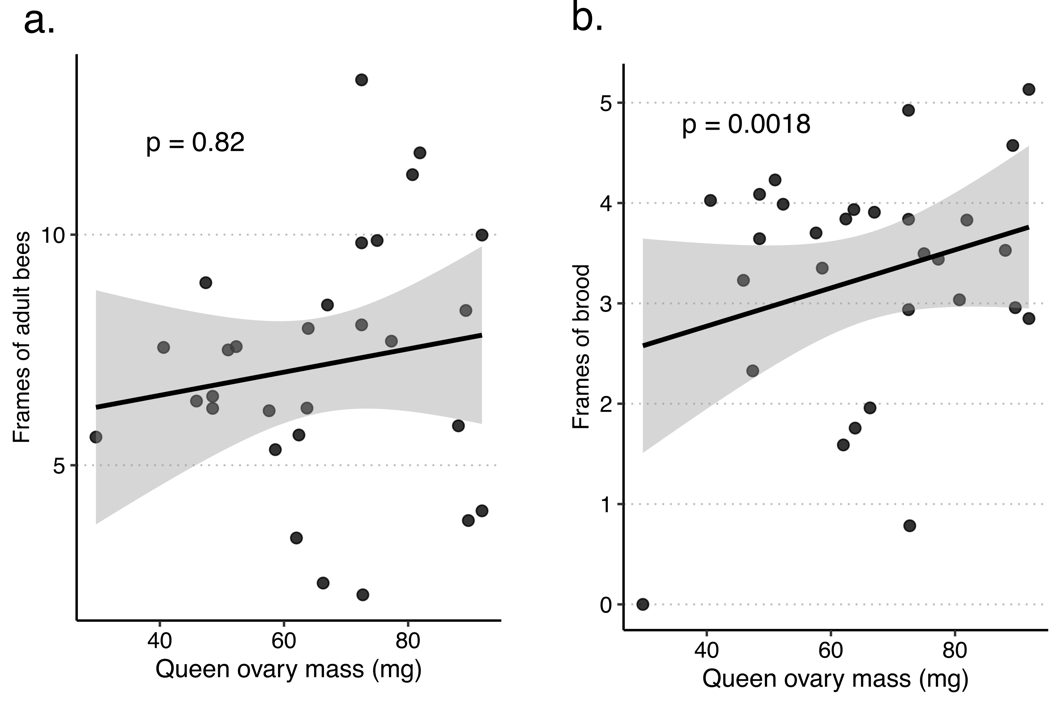

**Supplementary Figure 2. Relationship between amounts of brood and adult bees and queen ovary mass in the field experiment. a.** Queen ovary mass is not associated with frames of adult bees present in the colony at the end of the experiment after controlling for site and the adult bee population at the start of the experiment (linear model with fixed effects ovary mass (est. = 0.0082, t = 0.23, p = 0.82), initial bees (est. = 1.47, t = 3.12, p = 0.0045), and site (est. = -0.51, t = -0.41, p = 0.68). **b.** Queen ovary mass is positively associated with the amount of brood present in the colony at the end of the experiment after controlling for the same factors (linear model with fixed effects ovary mass (est. = 0.051, t = 3.50, p = 0.0018), initial bees (est. = 0.52, t = 2.64, p = 0.014), and site (est. = 1.73, t = 3.39, p = 0.0032).
