## Supplementary material for "Common viral infections inhibit egg laying in honey bee queens and are linked to premature supersedure": Supplmental Data 2 - qPCR sequences

**Primers for qPCR**

| Primer name | Bases | Sequence | Tm (50mM NaCl) C |
| --- | --- | --- | --- |
| BQCV-F | 17 | CGA AGC GTT TTC CGT GG | 54.4941 |
| BQCV-R | 19 | GCT GTC GAG AGT CAG AGT T | 53.92753 |
| DWVA-F | 28 | GTC TTG TGG ATG AAG GTT ATA TAA CTG G | 55.22766 |
| DWVA-R | 19 | TCC GTA GAA AGC CGA GTT G | 54.53572 |
| DWVB-F | 20 | ACC AAC GCG TGT CGT TCC TG | 60.54609 |
| DWVB-R | 20 | ACA AGT GGT TGG TCC CGT CG | 60.00336 |
| SBV-F | 23 | TAC GAA TCG TGA TTC GAT TCA TT | 52.3294 |
| SBV-R | 16 | ACG GGT CTG ACG CAA C | 55.33702 |

**Plasmid for qPCR standard curves**

Empty plasmid (pUC57) supplied by GenScript (https://www.genscript.com/vector/SD1176-pUC57_plasmid_DNA.html)

Insert sequence:

AAATTTGCTGACATCTAGTATGGTTCCCTCCGAAGTTTCCCTCAGGATAGCTGGCACTCGACCGTTTCTCATCGAACGCGTACGAGTCTCATCTGGTAAAGCGAATGATTAGAGGCCTTGGGGCCGAAACGACCTCAACCTATTCTCAAACTTTAAATGGGTGAGATCTCTGGCTTGCAAATTTGCTGACATCTAGTACGAAGCGTTTTCCGTGGGCTAAAGTGGTTGGTTTGGTGTACTCACCTCTTACTACGACCATTCCCGTAGATTATATAGTATACGGTCATTTTGAGGACGTGGAGTTGGGTTGTCCAACTTCAGGAATGTTGGCTCAAGCAGGTCTTAAAGTGCAACCTCCTACGAACTCTGACTCTCGACAGCAAATTTGCTGACATCTAGTAGCGCTAATACCAAGACACCAATCACGGACCTCACAAACACCACAGATGCTCAGGGTCGAGACTATATGTCTTACAAGAGCAGTCTTGTGGATGAAGGTTATATAACTGGAAAACAGAAGAAATATATAGCTATGTGGTGTAGTAAGCGTCGTGAACATACTGCTGACTTTGATCTTGTGTGGACTGATAATTTGCGTGTGTTAAGTGCGTATGTGCATGAACGTTCATCTTCAACTCGGCTTTCTACGGACTATATGTCTTACAAGAGCAACCAACGCGTGTCGTTCCTGATTGGACAACTGGTATTCTTGATATGGGTACCTTAAATATTCGTGTAATTGCTCCACTACGTATGAGTGCGACGGGACCAACCACTTGTCTATATGTCTTACAAGAGCAtacgaatcgtgattcgattcattatttcgcctgagtaaatcgagatttaccttgacggggtttgcttgacgttgcgtcagacccgtCTATATGTCTTACAAGAGCAgccctccataataagagtgtccacgtcgcggagggtgggactgaagcttttataatctttggtgcgaatcttataagtcttaccacatcttctggctgctggcatggtaagtagagcaggcaatgctcttgggatgagagatccattgccggatgttaaggtaatggcttaacaaggctgtgacgggtaacggtattactttgtaatattccggagaaggagcctgagagacggctactaagtctaaggattgtgcagcaggggcgaaacttgacctatggattttatctgaggcagtCTATATGTCTTACAAGAGCATTGGCTGGCCGTGATTTGACTGACTACCTCATGAAGATCCTTACAGAAAGAGGATATTCTTTCACTACTACGGCCGAACGTGAAATTGTTCGTGACATTAAAGAAAAACTCTGTTACGTTGCACTTGACTTTGAACAGGAAATGGCAACTGCTGCATCATCCTCAAGCTTGGAAAAGAGCTATGAACTTCCAGATGGTCAAGTAATTACTATTGGTAACGAAAGATTCCGTTGTCCCGAGGCTCTTTGTCGGTTATATTCTATATGTCTTACAAGAGCAgatgtttctccgttacgacgagtaaatcaggctatttggttgttatgtacaggtgctagggaagccgcgttccgtaatattaaaaccatcgccgaatgtttagccgatgaactcCTATATGTCTTACAAGAGCAgattcccgattggtttttgaataggcaaaaagatattgtagatggtaaatattcacagctcactagttcttatttggattcaaaacttcgtgaagatttggaacgaatgaagaaaattcgtgctcatagaggtttgcgtcattattggggttCTATATGTCTTACAAGAGCAacccaaactggaacgggacctgccttcttgaccatcaaggaatggatcgaaagagggactaccaaaagcatggaggcagcgaacatcatgagcaagcttccgaagactgttcgcacgccgactgacagctacatcagatccttcttcgaactgttacaaaatcctaaagtgagcaacgagcaattcctcaacaccgccgccacCTATATGTCTTACAAGAGCA**CAGTGCCTTACATTGCCAGTAGACCTTGGTTATATTGCATACGCCCTGAATCATCTTGGTTGAGTAAAGATAACAAGGATGGGGCTTTGATGTATAATTGTGTTAGTGGAATAGTGCGAGTAGAAGTT**CTATATGTCTTACAAGAGCA

Key:

Apo28S

BQCV

IAPV

DWVA

DWVB

SBV

Napis

AmActin

AmRPS5

AmRPS18

AmVg

**CrPV**
